# Feature-Specific Neural Reactivation during Episodic Memory

**DOI:** 10.1101/622837

**Authors:** Michael B. Bone, Fahad Ahmad, Bradley R. Buchsbaum

## Abstract

When recalling an experience of the past, many of the component features of the original episode may be, to a greater or lesser extent, reconstructed in the mind’s eye. There is strong evidence that the pattern of neural activity that occurred during an initial perceptual experience is recreated during episodic recall (neural reactivation), and that the degree of reactivation is correlated with the subjective vividness of the memory. However, while we know that reactivation occurs during episodic recall, we have lacked a way of precisely characterizing the contents—in terms of its featural constituents—of a reactivated memory. Here we present a novel approach, feature-specific informational connectivity (FSIC), that leverages hierarchical representations of image stimuli derived from a deep convolutional neural network to decode neural reactivation in *f*MRI data collected while participants performed an episodic recall task. We show that neural reactivation associated with low-level visual features (e.g. edges), high-level visual features (e.g. facial features), and semantic features (e.g. “terrier”) occur throughout the dorsal and ventral visual streams and extend into the frontal cortex. Moreover, we show that reactivation of both low- and high-level visual features correlate with the vividness of the memory, whereas only reactivation of low-level features correlates with recognition accuracy when the lure and target images are semantically similar. In addition to demonstrating the utility of FSIC for mapping feature-specific reactivation, these findings resolve the relative contributions of low- and high-level features to the vividness of visual memories, clarify the role of the frontal cortex during episodic recall, and challenge a strict interpretation the posterior-to-anterior visual hierarchy.

## Introduction

Not all our conscious memories for past events have the same quality of experience: some are vague and fuzzy, while others are sharp and detailed—sometimes nearly on par with the “fidelity” of direct perceptual experience. What accounts for this variability in the sharpness and “resolution” of memories? Researchers studying mental imagery, episodic memory, and working memory have over the last several decades or so converged on the idea that memories are constructed from the same neural representations that underlie direct perception (e.g. Ishai et al., 2002; Slotnick, Thompson, & Kosslyn, 2005; Polyn et al., 2005; Buchsbaum et al., 2012; Johnson & Johnson, 2014; Naselaris et al., 2015; Cabeza, Ritchey & Wing, 2015), a process known as neural reactivation (Danker & Anderson, 2010; Rissman & Wagner, 2012). Researchers have consistently reported that measures of neural reactivation throughout the dorsal and ventral visual streams reflect the content of episodic memory (Buchsbaum et al., 2012; Kuhl, Bainbridge, & Chun, 2012; Johnson & Johnson, 2014; St-Laurent et al., 2014; Cabeza, Ritchey & Wing, 2015), including low-level image properties such as edge orientation and luminosity (Harrison & Tong, 2009; Albers et al., 2013; Naselaris et al., 2015), as well as high-level semantic properties (Reddy, Tsuchiya & Serre, 2010; Cichy, Heinzle & Haynes, 2011). Moreover, the degree of neural reactivation has been demonstrated to correlate with the memory vividness (Cui et al., 2007; Johnson et al., 2015; St-Laurent, Abdi, & Buchsbaum, 2015; Dijkstra, Bosch & van Gerven, 2017; Bone et al., 2019).

The parallels between perception and memory extend beyond the representational overlap within posterior visual regions. As with perception, visual memory is subject to capacity constraints (Hesslow, 2012), necessitating the engagement of similar executive processes, such as selective attention (Hebb, 1968; Buschman & Miller, 2007; Johansson et al., 2012; Wynn et al., 2016; Bone et al., 2019) and working memory (Baddeley, 1988; Keogh & Pearson, 2014; Pearson, Naselaris, Holmes & Kosslyn, 2015). These executive processes serve to enhance and maintain neural reactivation of task-relevant image features within posterior visual regions via top-own projections from the frontal cortex (Mechelli, Price, Friston & Ishai, 2004; Nobre et al., 2004; Higo et al., 2011; Lee & D’Esposito, 2012; Dentico et al., 2014; Dijkstra, Zeidman, Ondobaka, Gerven, & Friston, 2017).

Although there is now strong evidence that a network of frontal cortical areas contributes to visual memory, there is currently a debate over the nature of the representations within these regions. According to one account, frontoparietal regions encode abstract task-level representations such as category membership (Freedman, Riesenhuber, Poggio, & Miller, 2001), rules (Warden & Miller, 2010; Riggall & Postle, 2012; Lee, Kravitz, & Baker, 2013), and stimulus-response mappings (Rowe, Hughes, Eckstein, & Owen, 2008). However, stimulus-specific responses have also been discovered within prefrontal regions (Miller, Erickson, & Desimone, 1996; Kuhl, Rissman, & Wagner, 2012; Ester, Sprague & Serences, 2015; St-Laurent, Abdi & Buchsbaum, 2015), with some subregions of the frontal cortex encoding both task-general and stimulus-specific representations in a high-dimensional state space (Mante, Sussillo, Shenoy, & Newsome, 2013; Rigotti et al., 2013; Raposo, Kaufman, & Churchland, 2014) to facilitate higher cognitive functions such as attention (Bichot, Heard, DeGennaro, & Desimone, 2015), working memory (Ester, Sprague & Serences, 2015) and decision making (Bizley, Jones, & Town, 2016). Whereas evidence for stimulus-specific representations within the frontal cortex has been growing rapidly over the last decade, there is still little information about the granularity of sensory features represented in frontal cortex, as the tools for detecting such representations are just beginning to emerge.

The detection of feature-specific neural representations has advanced significantly over the past few years with the advent of brain-inspired deep convolutional neural networks (CNN) (LeCun, Bengio, & Hinton, 2015). Early attempts at identifying and localizing neural activity associated with specific visual features focused on either high-level sematic/categorical features (Hung, Kreiman, Poggio, & DiCarlo, 2005; Meyers et al., 2008; Walther, Caddigan, Fei-Fei, & Beck, 2009; Reddy, Tsuchiya, & Serre, 2010; Smith, & Goodale, 2013) or low-level features such as edges (Kay, Naselaris, Prenger, & Gallant, 2008; Naselaris et al., 2015)—limiting findings to a small slice of the cortical visual hierarchy. In contrast, features extracted from the layers of a deep CNN have been linked to activity over nearly the entire visual cortex during perception, with a correspondence between the hierarchical structures of the CNN and cortex (Yamins et al., 2014; Güçlü and van Gerven, 2015; Wen et al., 2017; Eickenberg, Gramfort, Varoquaux, & Thirion, 2017; Seeliger et al., 2018).

Horikawa and Kamitani (2017) used this approach to reveal feature-specific neural reactivation throughout the ventral visual stream during mental imagery. However, because Horikawa and Kamitani were predicting category-average features, as opposed to image-specific features, the study was insensitive to the reactivation of lower-level features due to the large within-category variability of lower-level features relative to higher-level features. Moreover, the authors’ decoding approach did not account for the inherent correlations between the feature-levels extracted from the CNN. The architecture of feedforward CNNs is designed such that features from higher layers of the network are composed of features from lower layers, resulting in strong inter-layer correlations. Thus, any method that does not control for these inter-layer correlations will be prone to false positives, i.e. falsely detecting reactivation of features from (nearly) all levels of the visual hierarchy when only a small subset of the feature-levels are present within a given brain region. Güçlü and van Gerven (2015) and Seeliger et al., (2018) developed a method to address this issue that first assigns the layer that best predicts a given voxel/source’s activity to that voxel/source, and then uses the proportion of voxel/sources assigned to each layer within an ROI to infer the feature-levels represented within that cortical region. This approach, however, may overlook feature-levels that are weakly represented within a given region, due to the simplifying assumption that only one feature level is represented per voxel/source, resulting in false negatives.

To overcome some of these previous imitations in identifying feature-specific reactivation during memory recall, we introduce feature-specific informational connectivity (FSIC), a novel measure that incorporates a voxel-wise modeling and decoding approach (Naselaris et al., 2015), coupled with a variant of informational connectivity (Coutanche & Thompson-Schill, 2013; Anzellotti & Coutanche, 2018). Unlike previous measures of feature-specific neural reactivation, our method takes advantage of trial-by-trial variability in the retrieval of episodic memories by measuring the synchronized shifts in reactivation across cortical regions. We demonstrate that this approach eliminates false positives by accounting for inter-layer correlations while retaining sensitivity to more weakly represented features.

We used FSIC to examine feature-specific reactivation across the neocortex during a task requiring subjects to recall and visualize complex naturalistic images. The experiment consisted of three video viewing runs (Fig. 1b), used to train the encoding models, and three sets of alternating encoding and retrieval runs (Fig. 1a). During the encoding runs participants memorized a sequence of color images while performing a 1-Back task. In the following retrieval runs, the participants’ recall and recognition memory of the images was assessed. Feature-specific neural reactivation was measured while participants visualized a cued image within a light-grey rectangle, followed by a memory vividness rating. The participants then judged whether they had seen the identical image during encoding, followed by a rating of their confidence in this response.

**Figure 1.**
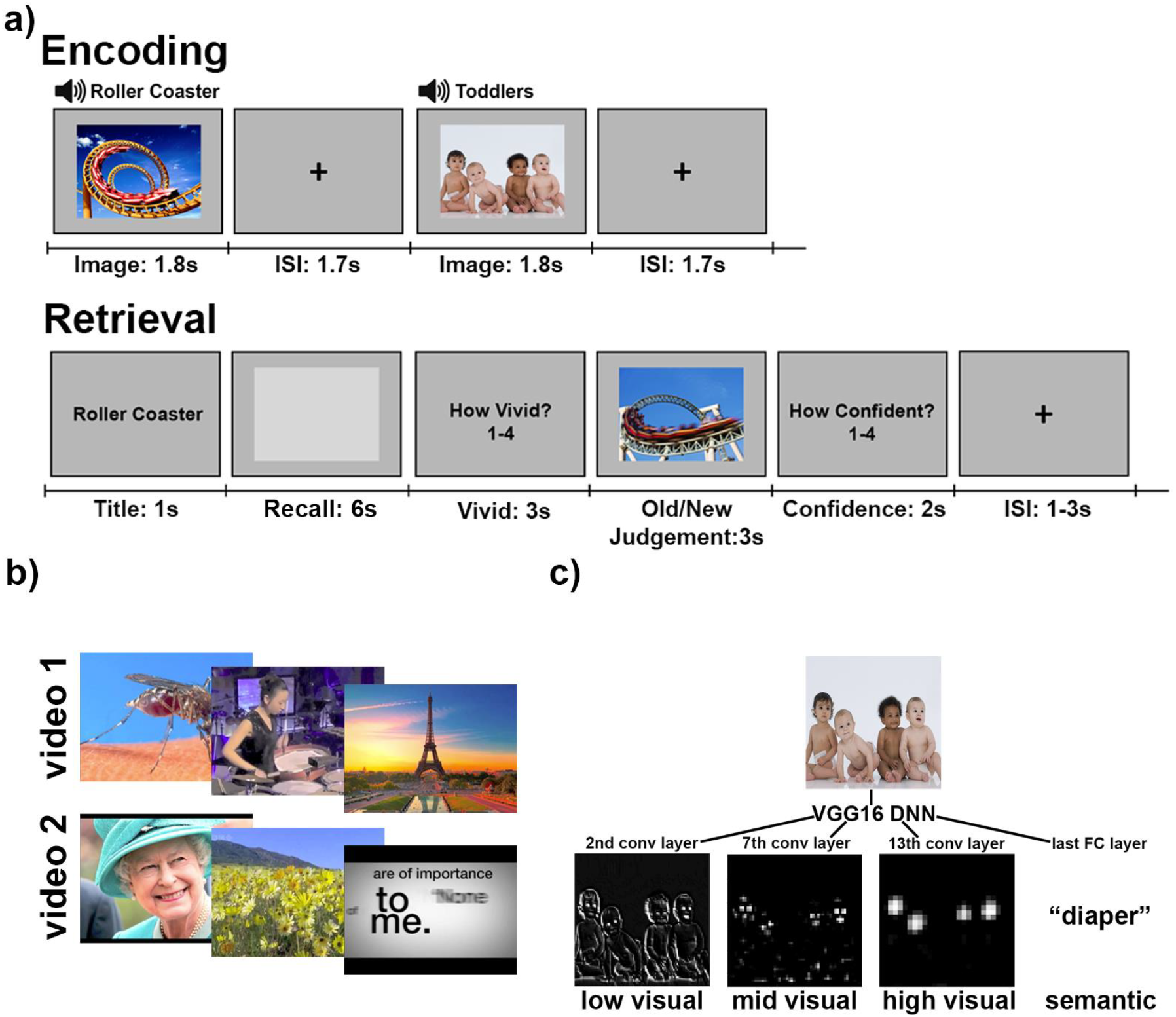
Procedure and Visual Features. a) Alternating image encoding and retrieval tasks. During encoding, participants performed a 1-back task while viewing a sequence of color photographs accompanied by matching auditory labels. During retrieval, participants 1) were cued with a visually-presented label, 2) retrieved and maintained a mental image of the associated photograph over a 6 second delay, 3) indicated the vividness of their mental image using 1-4 scale, 4) decided whether a probe image matched the cued item, and 5) entered their confidence rating with respect to the old/new judgement. b) Example stills from the two videos shown before the encoding and retrieval tasks. Data from the videos, which comprised a series of short clips, was used for training the encoding models. c) For each image, features were extracted from layer node activations using the VGG16 deep neural net (DNN). Activations from the 2^nd^, 7^th^ and 13^th^ convolutional layers, and the last fully connected layer were used, corresponding to low-visual, middle-visual, high-visual and semantic features, respectively.

Given the purported role of the frontal cortex in coordinating visual representations within posterior sensory regions (Mechelli, Price, Friston & Ishai, 2004; Nobre et al., 2004; Higo et al., 2011; Lee & D’Esposito, 2012; Dentico et al., 2014; Dijkstra, Zeidman, Ondobaka, Gerven, & Friston, 2017), we hypothesized that neural reactivation associated with all visual feature-levels should occur within—and be synchronized between—these cortical regions. Beyond establishing the cortical distribution of feature-specific visual representations, we were also interested in their connection to memory performance. To this end, we investigated the relationship between feature-specific reactivation during recall and both subjective (vividness ratings) and objective (recognition accuracy) behavioral memory measures. We hypothesized that reactivation of all feature levels would correlate with the vividness of the recalled image, but that lower level representations would have the strongest correlation because these features are most clearly associated with the sharp and intense phenomenology of vivid memories (Hebb, 1968; Kosslyn, Ganis, & Thompson, 2001). For the relation between reactivation and recognition memory, we hypothesized that high-level features should not assist in differentiating the encoded image and the lure due to the close semantic overlap between the two images (see Supplementary Figure 5 for example image pairs); thus, we hypothesized that only the reactivation of lower-level features during recall should correlate with recognition accuracy.

## Results

### Behavioral

#### Imagery Vividness Ratings

The mean vividness rating over trials, averaged across participants, was 3.04 (SD = 0.35). On average, 2.9% of trials were rated as vividness = 1, 19.7% as vividness = 2, 48.0% as vividness = 3 and 29.4% as vividness = 4. Participants failed to respond within the three second vividness rating period on 0.9% (SD = 2.2%) of the trials. These trials were excluded from all analyses.

#### Old/New Task Accuracy and Confidence Ratings

The means for old/new task accuracy and confidence ratings, averaged across participants, were 81.0% (SD = 11.0%; chance = 50%) and 3.46 (SD = 0.30), respectively. On average, 3.1% of trials were rated as confidence = 1, 10.9% as confidence = 2, 22.1% as confidence = 3 and 63.9% as confidence = 4. The association between accuracy and confidence ratings was significant (β = .89, p < .001) (measured using a generalized linear mixed-effects (LME) model with subject and image as crossed random effects). Participants failed to respond within the three second old/new response period on 1.0% (SD = 1.5%) of the trials, and the two second confidence rating period on 1.8% (SD = 2.3%) of the trials. The former trials were classified as incorrect, while the latter were excluded from analyses that incorporated confidence ratings.

### Neural Reactivation During Episodic Memory Recall

#### Measuring Neural Reactivation Using an Encoding-Decoding Approach

To measure neural reactivation during memory recall, an encoding-decoding approach was used (Naselaris et al., 2015). In short, encoding models were used to predict the expected neural activity in response to a set of features comprising a seen or imagined image. The correlation between model predictions and the activity measured during visual recall was then used to decode the cued image.

Brain activity measured during the encoding runs and the first two video runs were used to train cortical surface-based vertex-wise encoding models for each of four visual feature levels: low-level visual features, mid-level visual features, high-level visual features and semantic features. Given recent work showing a correspondence between visual features derived from an image recognition CNN and the features underlying human vision (Güçlü & van Gerven, 2015; Horikawa & Kamitani, 2017), the encoding models used features extracted from layer activations in a DNN (VGG16; Simonyan & Zisserman, 2014) to predict neural activity (Fig. 1c). Based on the findings of Güçlü & van Gerven (2015), outputs from the units in the second, seventh, thirteenth, and sixteenth layers of VGG16 were used as approximations of low-visual, mid-visual, high-visual and semantic cortical features, respectively.

To identify brain regions that were accurately characterized by the vertex-wise feature-specific encoding models, neural activity predicted by the encoding models for each trial and feature-level were grouped into 148 bilateral cortical Freesurfer ROIs (Destrieux, Fischl, Dale, & Halgren, 2010). For each ROI and trial, predictions of neural activity for all encoded images were generated and correlated with the observed neural activity. The predictions were then sorted by correlation coefficient, and the rank of the prediction associated with the actual cued image was recorded. To make the rank measure more interpretable, the rank was subtracted from the mean rank so that a value significantly greater than 0 indicates neural reactivation (i.e. the cued image could be decoded from neural activity during recall).

Figure 2 depicts neural reactivation for all cortical ROIs during episodic recall. Consistent with previous findings (Buchsbaum et al., 2012; Johnson & Johnson, 2014; St-Laurent et al. 2014; Cabeza, Ritchey & Wing 2015; Horikawa & Kamitani, 2017), the ability to decode recalled memories was greatest throughout the dorsal and ventral visual streams for all feature levels, with the peak ROIs (Figure 4a; see ROI/Seed Selection for details) located within brain regions of the cortical visual processing hierarchy associated with each feature level. Significant decoding accuracy was also seen in the lateral prefrontal cortex, particularly within the inferior frontal sulcus. Overall, our findings indicate widespread neural reactivation associated with all feature-levels during episodic recall.

**Figure 2.**
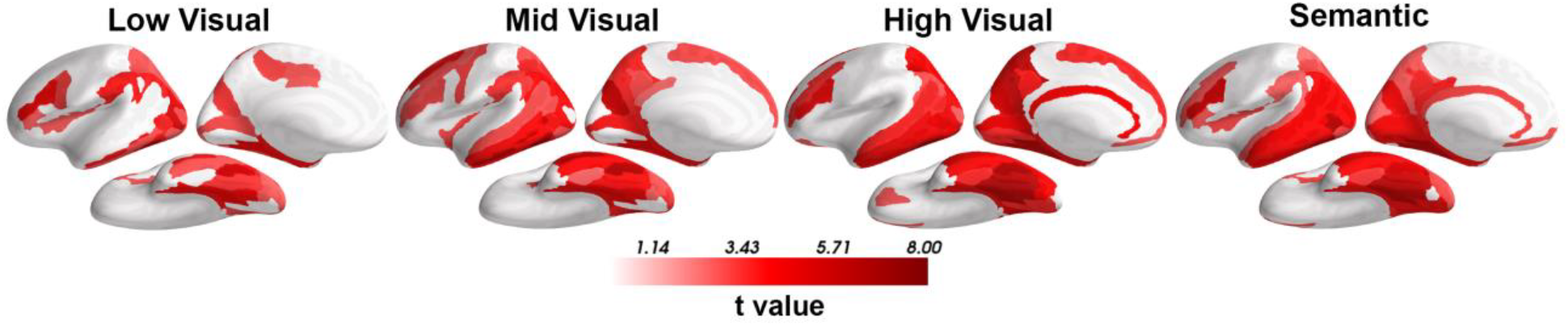
Neural Reactivation During Episodic Recall. Reactivation for each bilateral ROI and feature level (column = feature). Reactivation was significantly greater than chance throughout the dorsal and ventral visual streams and within the lateral and orbital frontal cortex during recall. The t-values are thresholded at p < 0.05, FDR corrected.

#### Feature-Specific Informational Connectivity

Despite strong findings indicating reinstatement of all CNN feature-levels throughout the cerebral cortex, correlations between features from different network layers (Supplementary Figure 1) makes it difficult to independently assess the contribution of each feature level to memory reactivation. For example, because higher level features are composed of lower level features, it may be the case that only neural activity associated with low-level visual features is reactivated in a given ROI, but, due to the correlation between high- and low-level visual features, activity associated with high-level visual features also appears to be reactivated. This could explain why reactivation of semantic features was found within the calcarine sulcus (Figure 2, last column), which is surprising given the area’s assumed role in low-level visual processing (Güçlü & van Gerven, 2015). Thus, to assess the independent contribution of each feature level to reactivation, it is necessary to statistically account for neural activity associated with all non-target features. To that end, we developed feature-specific informational connectivity (FSIC)—a variant of informational connectivity (Coutanche & Thompson-Schill, 2013). FSIC measures the correlation of feature-specific neural reactivation between a seed ROI and a target ROI, while covarying out all non-target feature-levels, thereby enabling the detection neural reactivation associated with a specific set of features.

Before applying FSIC to experimental data we first validated the approach with a simulation to determine whether FSIC can detect neural reactivation associated with a specific visual feature-level, while eliminating false positives. To this end, fMRI data was simulated for 200 subjects using the node activations/outputs from the CNN in response to the experimental stimuli (see fMRI Data Simulation for details). Figure 3a depicts the classification accuracy results for this simulated data. Despite each ROI representing features from only one feature-level, significant effects are present for all feature-levels within each ROI. If classification accuracy were to be used to infer the representation of features within a given ROI, it would lead to the false conclusion that each ROI contains representations of all feature-levels. Figure 3b depicts neural reactivation results using FSIC assuming identical trial-by-trial reactivation fidelity (i.e. proportion of forgotten features) across feature-levels. In contrast to the naïve classification accuracy method, FSIC accurately identifies neural reactivation associated with only the features present within each ROI, albeit with a small amount of signal smearing to features in adjacent layers. No signal smearing was found when trial-by-trial reactivation fidelity was assumed to be independent across feature-levels (see diagonal of Supplementary Figure 2c)—an assumption that more accurately modeled the off-diagonal of figure 4b (compare Supplementary Figures 2b and 2c)—so figure 3b likely overestimates the amount of signal smearing one can expect when applying the technique to real data. Moreover, similar, yet generally weaker, results were found when the seed ROIs contained an equal proportion of voxels representing each feature-level (seeds in the above results only contained the target feature-level), suggesting that the feature-specificity of FSIC is not dependent on the selection of seed ROIs that only contain the target feature-level (Supplementary Figure 2a). FSIC may therefore be used to greatly improve our ability to isolate neural reactivation associated with specific features when compared to the naïve approach.

**Figure 3.**
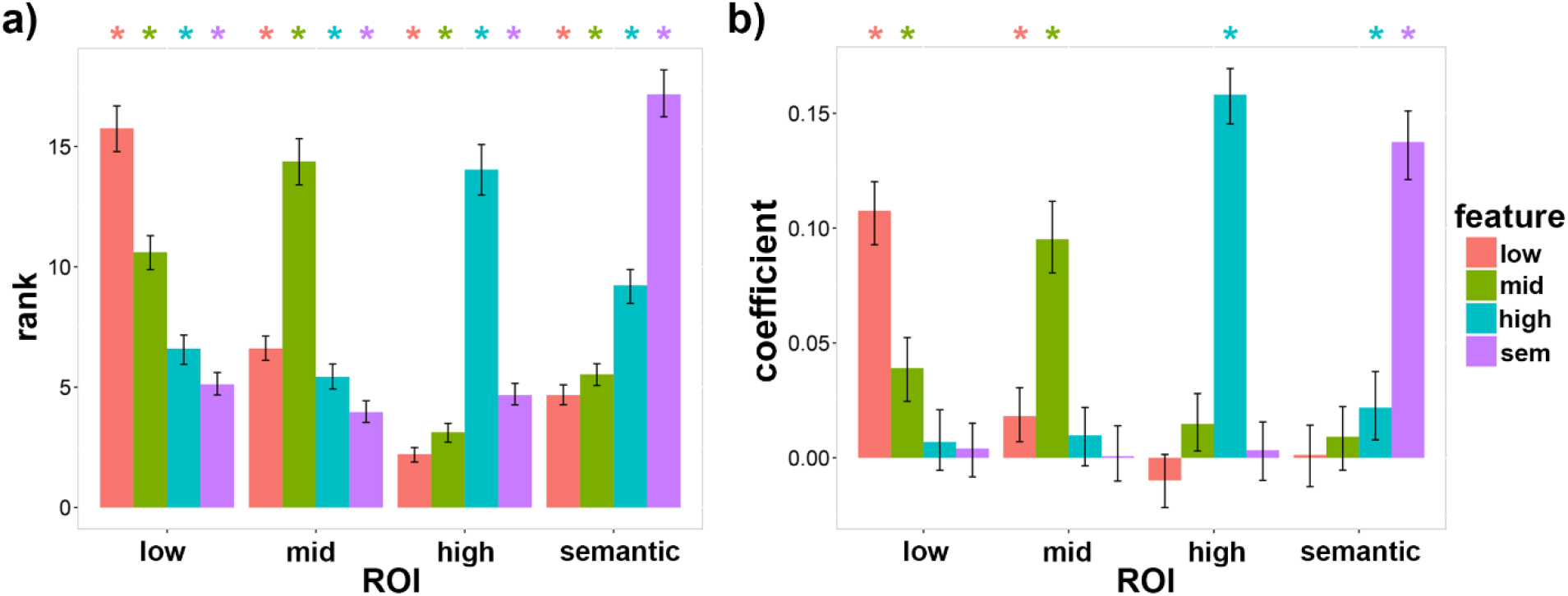
Simulated Results for Decoding Accuracy and Feature-Specific Informational Connectivity. fMRI data was simulated (200 simulated subjects; see Methods section) and then run through the processing pipeline for FSIC (see Methods section) to validate the approach. ROIs only contain features from the indicated feature-level. a) Image classification performance (rank measure) for all combinations of ROI and feature-level. Correlations between feature-levels result in the classification accuracy measure falsely indicating the presence of features that are not present within the target ROI. b) FSIC results for all combinations of ROI and feature-level assuming identical trial-by-trial memory accuracy across feature-levels. A separate seed was used for each feature-level corresponding to that feature-level (the results are also depicted in Supplementary Figure 2b along the diagonal). Significant FSIC results only indicate the presence of the feature-level contained within each ROI, except for relatively weak evidence for the presence of adjacent feature-levels (e.g. a significant effect associated with mid-level features was found within the low-level ROI). Error bars are 90% CIs; * indicates p < 0.05, FDR corrected.

**Figure 4.**
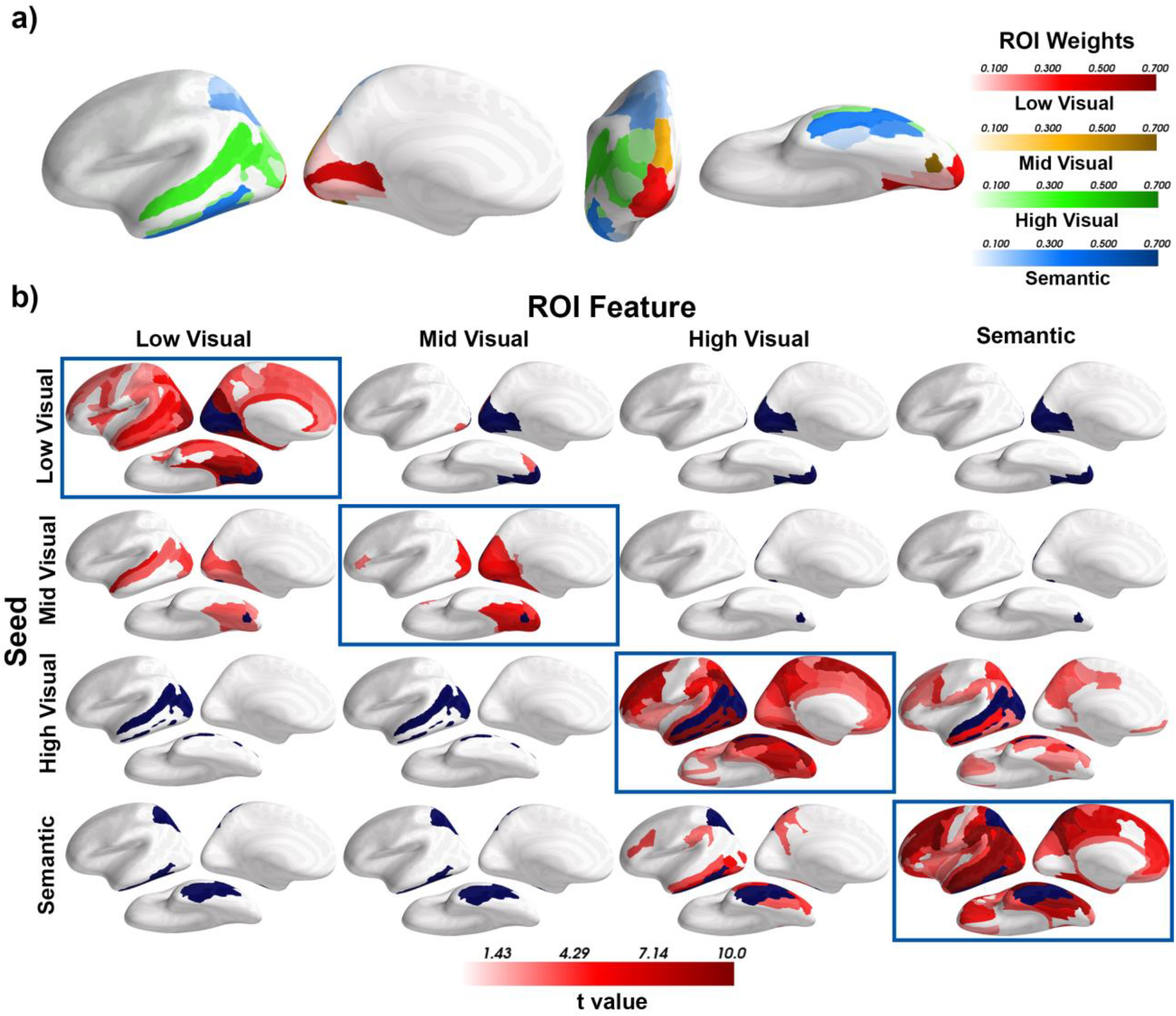
Seed ROI Weights and Feature-Specific Informational Connectivity during Episodic Recall. All ROIs are bilateral. a) Seed ROI weights for each feature level. Seed ROI weights are proportional to the decoding accuracy for the target feature level relative to the other feature levels during encoding/perception (i.e. the relative accuracy peaks). b) FSIC results for all combinations of seed ROI/feature-level and target ROI feature-level. For FSIC, neural reactivation (during memory recall) of each feature level within the corresponding seed ROI (rows; seed ROIs colored dark blue) was correlated with reactivation of all four feature levels (columns), controlling for the reactivation of the non-target feature levels, within all ROIs except for the seed. The t-values associated with those correlations are indicated with shades of red and thresholded at p < 0.05, FDR corrected.

Figure 4b depicts the results obtained from applying FSIC to fMRI data measured during visual recall (i.e. the task depicted in Figure 1a). Specifically, the figure displays the correlation of neural reactivation for each feature level within the corresponding seed ROIs (rows; ROIs from Figure 4a marked with blue) and all four target feature levels within all other ROIs (columns), controlling for all non-target feature levels. Off-diagonal results indicate the correlation between different feature-levels, whereas on-diagonal results indicate the correlation within the same feature-level (Figure 3 depicts a simulation of the latter). According to our simulation results, the generally weak correlations within the off-diagonal indicate that trial-by-trial variation in memory reactivation is largely independent across feature-levels (Supplementary Figures 2b-c), i.e. the degree of reactivation from one feature-level is only weakly related to the amount of feature reactivation from a different feature-level. In contrast, strong correlations along the diagonal indicate widespread neural reactivation for low-visual, high-visual, and semantic features that extends beyond the occipital cortex into higher-order regions of the dorsal and ventral visual streams as well as the frontal cortex. Reactivation of mid-level features was, however, primarily limited to the occipital cortex; and this difference is not due to the relativity small size of the mid-level seed ROI (see Supplementary Figure 3). Although expected for higher-order features (Freedman, Riesenhuber, Poggio, & Miller, 2001; Wagner, Paré-Blagoev, Clark, & Poldrack, 2001; Huth, Nishimoto, Vu, & Gallant, 2012; Carota, Kriegeskorte, Nili, & Pulvermüller, 2017), the widespread presence of low-level visual features within higher-order regions (see Table 1) appears to challenge a strict interpretation of the cortical visual hierarchy, which would predict results similar to what we observed for mid-level visual features.

**Table 1.** Low-Level Feature-Specific Informational Connectivity During Imagery within the Frontal Cortex. The table lists the significant FSIC results (and associated statistics) within the frontal cortex depicted in the first row and first column of figure 4b.

| Frontal ROI | $\beta$ | SE | t value | Lower bound | Upper bound | P (FDR corrected) |
| --- | --- | --- | --- | --- | --- | --- |
| Middle frontal sulcus | 0.083 | 0.021 | 3.90 | 0.050 | 0.117 | 0.004 ** |
| Superior precentral sulcus | 0.083 | 0.021 | 3.90 | 0.047 | 0.117 | 0.004 ** |
| Superior circular sulcus | 0.083 | 0.021 | 3.89 | 0.047 | 0.115 | 0.004 ** |
| Inferior precentral sulcus | 0.083 | 0.021 | 3.87 | 0.046 | 0.118 | 0.004 ** |
| Superior frontal gyrus | 0.077 | 0.022 | 3.47 | 0.039 | 0.114 | 0.004 ** |
| Anterior midcingulate | 0.068 | 0.022 | 3.16 | 0.034 | 0.104 | 0.004 ** |
| Superior frontal sulcus | 0.067 | 0.023 | 2.97 | 0.027 | 0.107 | 0.008 ** |
| Middle frontal gyrus | 0.057 | 0.022 | 2.61 | 0.022 | 0.093 | 0.010 * |
| Anterior cingulate | 0.055 | 0.022 | 2.55 | 0.018 | 0.091 | 0.023 * |
| Short Insular gyri | 0.053 | 0.021 | 2.48 | 0.016 | 0.090 | 0.022 * |
| Precentral gyrus | 0.053 | 0.022 | 2.44 | 0.018 | 0.089 | 0.016 * |
| Inferior frontal gyrus - opercular | 0.048 | 0.022 | 2.25 | 0.013 | 0.083 | 0.020 * |

### Neural Reactivation and Behavioral Performance

#### Correlation Between Neural Reactivation and Vividness Ratings

With the cortical distribution of feature-specific neural reactivation established, we then assessed how feature-specific reactivation relates to behavioral measures of memory performance. To test the hypothesis that memory vividness is largely the result of the reactivation of lower-level visual features (Hebb, 1968; Kosslyn, Ganis, & Thompson, 2001), measures of low- and mid-level reactivation (lower-level features), and high-level and semantic reactivation (higher-level features) were combined (averaged) together, along with the associated ROIs (Figure 5a), forming four separate reactivation measures: one for each unique combination of feature-level and ROI. The within-subject correlations between these reactivation measures and vividness was then assessed using a LME model, with the vividness rating as the dependent variable (DV), the four reactivation measures as independent variables (IV), and the subject and image as crossed random effects (random-intercept only, due to model complexity limitations), thereby controlling for the correlations between feature-levels and ROIs. Figure 5b depicts the partial regression coefficients associated with the four reactivation measures. Contrary to our hypothesis, reactivation of lower-level and higher-level features, within corresponding ROIs, appear to contribute approximately equally to subjective vividness.

**Figure 5.**
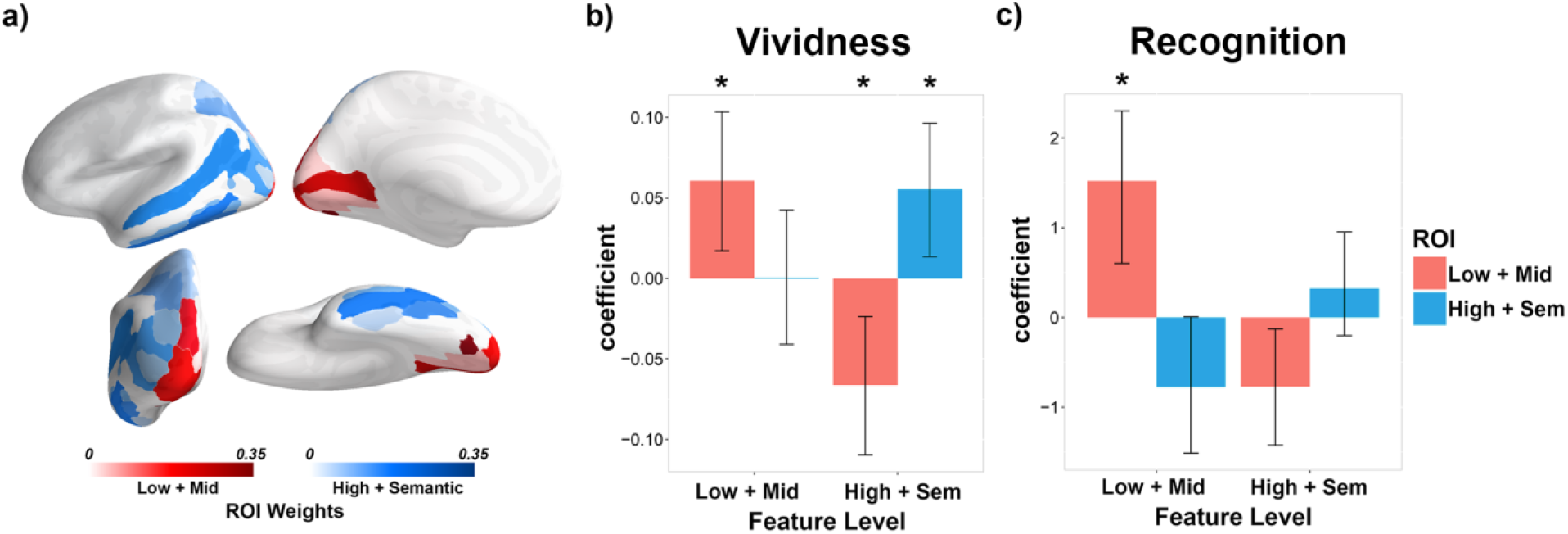
Correlations Between Feature-Specific Neural Reactivation, Vividness and Recognition Accuracy. a) ROI weights combining the low- and mid-level and high- and semantic-level ROIs. b) Within-subject partial regression coefficients measuring the relation between neural reactivation during recall and vividness for all combinations of feature-level and ROI. c) Between-subject partial regression coefficients measuring the relation between neural reactivation and recognition accuracy (during the Old/New task) for all combinations of feature-level and ROI. The error bars are 95% CI; * indicates p(β = 0) < .05, FDR corrected.

In addition to the positive partial correlations, we found that reactivation of higher-level features within the lower-level ROI negatively correlated with vividness. According to the predictive coding account of perception, top-down connections from neurons that encode high-level/semantic features drive neural activity representing lower-level features to generate a model of the expected stimulus, which is compared against the perceptual input to generate an error signal (Rao & Ballard, 1999; Friston, 2005, 2010; Bastos et al., 2012). From this perspective, the presence of higher-level features within the lower-level ROI may represent the top-down inference of lower-level features. When reactivation of the perceived low-level features is statistically controlled for—as in the above analysis—the reactivation of higher-level features within the lower-level ROI is constrained to represent only the *incorrect* inferences, i.e. predictions of low-level features that were not present in the encoded image. These incorrect inferences would result in mental imagery of a generic image associated with the recalled high-level/semantic features which participants were instructed not to rate as vivid—even if the generic mental image contained many visual details. Therefore, the observed negative partial correlation between vividness and neural reactivation of higher-level features within the lower-level ROI is consistent with a predictive coding account of perception and memory recall.

#### Correlation Between Neural Reactivation and Recognition Accuracy

We hypothesized that recognition accuracy during the old/new task would selectively correlate with reactivation associated with lower-level visual features during episodic memory recall, due to the lure image being semantically similar to the original image but differing with respect to its low-level visual features. To test this claim, the same analytical approach described above for the correlation between reactivation and vividness was used, replacing vividness with accuracy as the DV (correct = 1, incorrect = 0). No significant correlations were found [Low Feature, Low ROI: β = .001, p = .972; Low Feature, High ROI: β = .037, p = .318; High Feature, Low ROI: β = -.010, p = .863; High Feature, High ROI: β = -.031, p = .318; two-tailed, FDR corrected]. Next, we examined the correlation between-subjects using a similar model to the one used for the within-subject analysis (except the DV and IVs were within-subject averages, and subject and image were not used as random effects). Consistent with our hypothesis, we found a significant partial correlation between recognition memory accuracy and lower-level reactivation within the corresponding ROI (Figure 5c; within- and between-subject coefficients were pooled together when controlling for multiple comparisons using FDR).

The between-subject result suggests that only some subjects may successfully use neural reactivation within early visual regions to enhance recognition memory accuracy. The null within-subject finding might therefore be a consequence of this individual difference. To explore this possibility, the correlation between memory accuracy and neural reactivation was calculated for each subject individually (using the within-subject linear model, except subject and image were not used as random effects). The resulting partial correlation coefficients for each combination of ROI and feature-level were then separately correlated with the subjects’ memory accuracy for all trials, lure trials, and ‘old’ trials (Supplementary Figures 4a-c, respectively). Significant positive correlations were found for lower- and higher-level features on lure trials, suggesting that the hypothesized positive within-subject correlation between memory accuracy and neural reactivation may only be evident for subjects with relatively high recognition accuracy. This possibility was tested using the same model as the original within-subject analysis on the thirteen subjects (half of the twenty-seven) with the highest average memory accuracy on the lure trials (Supplementary Figure 4d; the thirteen subjects with the lowest average memory accuracy were also tested: Supplementary Figure 4e). For this high-performance group, lower-level features within the corresponding ROI positively correlated with memory accuracy [β = .081, p = .014]. Thus, there is a relationship between low-level feature reactivation and recognition memory performance, but it is limited to the higher performing subset of participants.

## Discussion

The primary goal of the current study was to identify the specific feature-level composition of neural reactivation patterns measured throughout the neocortical mantle during a task requiring vivid recall of a diverse set of naturalistic images. We found that neural reactivation of low-level, mid-level, high-level, and semantic features occurs throughout the cortex, including much of the dorsal and ventral visual streams and frontal cortex. A classic theory relating neural reactivation to the subjective qualities mental imagery postulates that the vividness of mental imagery is primarily based upon neural reactivation of lower-level visual features, e.g. edges and simple shapes (Hebb, 1968). We found, however, that reactivation of lower- and higher-level features contribute approximately equally to subjective vividness. We further hypothesized that distinguishing two semantically and visually similar images during a visual recognition memory task would benefit from neural reactivation of lower-level visual features, particularly within early visual regions. Consistent with our hypothesis, subjects with greater lower-level reactivation within the early visual cortex during recall had greater recognition accuracy. Moreover, in a within-subject analysis, we showed that trial-to-trial variation in low-level feature reactivation predicted greater recognition accuracy, albeit only for participants with higher-than-average recognition performance on lure trials.

### Feature-Specific Neural Reactivation during Episodic Memory

Using FSIC, we found that visual features from all selected levels of the CNN were represented, to some degree, throughout the cortical visual hierarchy; but these representations were not evenly distributed across ROIs (see the diagonal of Figure 4b). Consistent with previous work indicating a correspondence between the hierarchical organization of the layers of a CNN and the cortical regions of the visual processing stream (Güçlü & van Gerven, 2015; Horikawa & Kamitani, 2017), the distribution of features, revealed by FSIC and the locations of peak neural reactivation for each feature-level (Figure 4a), was organized according to the posterior-to-anterior cortical visual hierarchy. This organization was also evident in the correlations between neural reactivation, imagery vividness ratings and recognition response accuracy (Figure 5), thereby establishing the ability to use CNN feature vectors to more precisely characterize and decompose the content of episodic memories.

Consistent with prior work we found that the association between feature-levels from the CNN and neural activity was largely congruent with the well-known hierarchical organization of the visual system. However, we also found strong evidence for lower-level visual features represented within higher-order cortical regions, and higher-level features within lower-order regions (Figure 4b). Unlike strictly feed-forward CNN’s, like the one used in this study, the cortex comprises a complex network of both feed-forward and feed-back connections that can bypass intermediate areas, facilitating direct communication between lower- and higher-order regions (Desimone et al., 1984; Lamme, Super & Spekreijse, 1998; Hegde & Felleman, 2007; Kravitz et al., 2013), thereby enabling the maintenance, modulation and combination of features at multiple levels (Chun & Jiang, 1999; Hopfinger, Buonocore & Mangun 2000; Gilbert & Sigman, 2007; Zanto, Rubens, Bollinger & Gazzaley, 2010; Zanto, Rubens, Thangavel, & Gazzaley, 2011; Gazzaley & Nobre, 2012; Piëch, Li, Reeke & Gilbert, 2013). For example, the inferior frontal gyrus has been implicated in the selective maintenance of task-relevant visual information via top-down connections with the visual cortex during working memory and mental imagery (Vandenberghe et al., 1996; Nobre et al., 2004; Mayer et al., 2007; Zanto, Rubens, Bollinger & Gazzaley, 2010; Higo et al., 2011; Dijkstra, Zeidman, Ondobaka, Gerven, & Friston, 2017). Through the use of FSIC, we show that the inferior frontal gyrus contains representations of visual features from all levels of the visual hierarchy during the recall of naturalistic scenes, and that the reinstatement of these representations within the IFG are correlated with the reinstatement of the same features within the occipital cortex, supporting the idea that the region facilitates feature-specific neural reactivation in early visual areas.

Low-level visual, high-level visual, and semantic features, but not mid-level visual features, were identified in many higher-order visual and frontal regions beyond the inferior frontal gyrus. While this was expected of high-level and semantic features (because the features represent object identity), the prevalence of low-level features within the frontoparietal cortex and higher-order regions of the ventral visual stream was more surprising. This raises the question of the function of such low-level features within these putative higher-order regions—a question that recent advances within the field of computational neural networks may shed some light upon. Like the receptive fields of neurons within the visual cortex (Smith, Singh, Williams, & Greenlee, 2001; Rolls, Aggelopoulos, & Zheng, 2003), the nodes that comprise feedforward CNNs designed to perform visual classification and localization tasks are organized such that the lower-order layers have small receptive fields and weak semantics, whereas the higher-order layers have large receptive fields and strong semantics (Luo, Li, Urtasun, & Zemel, 2016). Consequently, the resolution of the semantic-sensitive layers is low, resulting in the loss of fine details essential for some tasks (e.g. the classification of small objects). To address this problem, more recent CNNs have incorporated top-down connections and “skip” connections (which bypass adjacent layers) to directly combine the outputs of lower- and higher-order layers of the network, thereby increasing the effective resolution of the semantic-sensitive layers (Liu et al., 2018). This approach has been proven to be effective for a variety of tasks requiring both accurate semantics and fine visual details, including classification and localization of small objects (Shrivastava, Sukthankar, Malik, & Gupta, 2016), and salient object detection (a key element of attentional processes) and boundary delineation (important for the coordination of grasping behavior, among other tasks) (Zhang et al., 2017). Given the functional roles of the higher-order ventral visual stream in visual object classification (Grill-Spector, & Weiner, 2014) and the frontoparietal cortex in attention and grasping behavior (Ptak, 2012; Ptak, Schnider, & Fellrath, 2017), we posit that the presence of low-level visual representations within these regions may likewise facilitate visual classification, attentional allocation and motor planning tasks specifically, and any task that requires both accurate semantics and fine visual details more generally.

### Feature-Specific Neural Reactivation and Memory Vividness

To investigate the functional contributions of feature-specific neural reactivation to memory, we tested the hypothesis that the vividness of memory recall should positively correlate with the degree of neural reactivation encoding visual features—particularly low-level visual features. Although previous research had found correlations between vividness and neural reactivation throughout early and late regions of the ventral and dorsal visual streams (Cui et al., 2007; Johnson et al., 2015; St-Laurent, Abdi, & Buchsbaum, 2015; Dijkstra, Bosch & van Gerven, 2017; Bone et al., 2019), the relative contributions of the reinstatement of lower- and higher-level visual features remained an open question. By measuring the reactivation of features from different levels of the visual hierarchy, as opposed to inferring feature-level based upon the location of reactivation (i.e. reverse inference (Poldrack, 2011)), we found that the reinstatement of lower- and higher-level visual features correlated with vividness to an approximately equal degree. While our hypothesis did predict that vividness should correlate with reinstatement of both low- and high-level visual features, the low-level correlation was expected to be stronger based upon the assumption that the recall of visual details constituting a vivid memory is primarily dependent upon the reinstatement of low-level features (Hebb, 1968; Kosslyn, Ganis, & Thompson, 2001).

This assumption, however, may overlook the inference of low-level features from high-level features. According to the predictive coding account of perception, visual experience results from the reciprocal exchange of bottom-up and top-down signals throughout the cortical hierarchy (Rao & Ballard, 1999; Friston, 2005, 2010; Bastos et al., 2012). Top-down signals from neurons representing high-level features, which encode statistically and behaviorally significant non-linear combinations of lower level features, serve to drive and/or modulate neural activity representing the associated lower-level features—thereby functioning as a generative model of how environmental stimuli cause sensations. During perception, these top-down connections convey predictions, which are compared against the perceptual input to generate an error signal. This signal is then propagated back up the hierarchy to update the predictions (i.e. alter higher-order activations) and enhance memory of the features that diverged from expectations (Axmacher et al., 2010; Henson & Gagnepain, 2010). During episodic memory recall, cued higher-level features are used to infer lower-level features, while the sparsely recalled lower level features that were not accurately predicted during perception serve to constrain this inference to be specific to the recalled episode. Therefore, according to a predictive coding account of visual recall, the number and accuracy of remembered visual details (i.e. memory vividness) should depend upon the reactivation of both high- and low-level features. Moreover, because participants were instructed to not rate generic imagery related to the cue as vivid, the top-down inference of low-level features that were not present in the encoded image should correlate negatively with vividness, which is what we found. Thus, the partial correlations between subjective vividness and feature-specific neural reinstatement are consistent with a predictive coding account of visual perception and memory recall.

### Feature-Specific Neural Reactivation and Recognition Accuracy

Whereas our vividness results serve to demonstrate a connection between feature-specific neural reactivation and the subjective quality of memory, we were also interested in establishing the relationship between neural reactivation and an objective memory measure: recognition memory accuracy. The recognition memory task participants performed for our study required access to fine-grained memory information in order to identify a probe image drawn from the same semantic category (e.g. two images of a steam train) as old or new. Given the strong semantic overlap between the two images, higher-level semantic-like features alone would be unlikely to provide enough information to distinguish the images. Consequently, we hypothesized that the recall of lower-level features would be required to perform well on the task. Overall, our results supported this hypothesis (Figure 5d and Supplementary Figure 4d). We found that reactivation of lower-level features within the early visual cortex positively correlated with recognition accuracy within- and between-subjects, albeit the within-subject result only held for subjects with greater-than-average recognition accuracy on lure trials.

What might be the cause of this individual difference in the relationship between neural reactivation and recall accuracy? One possibility is that the participants differ in their reliance upon the reinstatement of higher-vs. lower-level features when comparing the presented image with the memorized image. Our original hypothesis that reactivation of lower-level features should positively correlate with recognition accuracy within-subjects assumed that *all* subjects would utilize lower-level representations when performing the task. Our failure to find the hypothesized within-subject effect appears to be the result of greater than expected individual variation in the ability or tendency of subjects to reactivate low-level visual features during memory retrieval. Future studies will be required to explore the cause and implications of these important individual differences.

### Limitations

We have demonstrated that features at all levels of the visual hierarchy are reactivated throughout the cortex during episodic recall. Moreover, we have shown how such feature-specific reactivation relates to the vividness of recall, and subsequent recognition accuracy. To obtain these findings, it was necessary to develop and utilize methods that control for the inherent correlations between feature-levels (e.g. FSIC). Although our simulation results strongly indicate that our methods were successful in this regard, a caveat must be considered. As with any model of feature-specific cortical representations, the features extracted from VGG16 (the CNN used in this study) cannot be expected to be a complete set of all visual features represented within a given participant’s cortical activity. Consequently, our approach cannot exhaustively control for all inter-level correlations, potentially resulting in the false detection of feature-specific neural reactivation. To address this concern, consider two hypotheticals: lower-level features tend to be detected within regions that only contain higher-level features, and/or higher-level features tend to be detected within regions that only contain lower-level features. If the former was true, we would expect approximately equal reactivation of mid-level features relative to low-level features within higher-order cortical regions, due to the mid-level and low-level features correlating with higher-level features to approximately the same degree (Supplementary Figure 1). In contrast, low-level reactivation was much more pronounced than mid-level reactivation (Figure 4b; first two rows along the diagonal). If the latter was true, then semantic features would be expected within the earliest region of the visual cortex: the calcarine sulcus. This was not the case (Figure 4b; last row along the diagonal). Therefore, our results strongly indicate that the methods used within the current study, e.g. FSIC, successfully controlled for the correlations between features drawn from different levels of the visual hierarchy, thereby eliminating the false positives that previous approaches (e.g. Horikawa & Kamitani, 2017) were susceptible to.

## Conclusion

The contributions of this study were fourfold. First, we developed novel measures of feature-specific neural reactivation, e.g. FSIC, that control for the inherent correlations between hierarchically organized feature-levels without sacrificing sensitivity. Second, the results obtained from FSIC revealed that neural reactivation during episodic memory is more widespread than previously thought—particularly for low-level features (e.g. edges)—which we posit subserves a multitude of cognitive functions requiring both fine visual detail and accurate object/scene categorization (e.g. fine grasping behavior). Third, we found that neural reactivation of lower-level and higher-level visual features contributed equally to the subjective vividness of recall, which we argue supports a predictive coding account of perception and recall. Lastly, we confirmed that when differentiating semantically nearly-identical images from memory only reactivation of low-level visual features correlates with recognition accuracy. Overall, the multifaceted nature of the current study’s results shows the potential for FSIC, and other feature-specific approaches to decomposing neural pattern representations, to test and elucidate the mechanisms underpinning long held theories about the brain basis of memory and cognition.

## Materials and Methods

### Participants

Thirty-seven right-handed young adults with normal or corrected-to-normal vision and no history of neurological or psychiatric disease were recruited through the Baycrest subject pool, tested and paid for their participation per a protocol approved by the Rotman Research Institute’s Ethics Board. Subjects were either native or fluent English speakers and had no contraindications for MRI. Data from ten of these participants were excluded from the final analyses for the following reasons: excessive head motion (5; removed if > 5mm within run maximum displacement in head motion), fell asleep (2), did not complete experiment (3). Thus, twenty-seven participants were included in the final analysis (15 males and 12 females, 20-32 years old [mean: 25]).

### Stimuli

111 colored photographs (800 by 600) were gathered from online sources. For each image, an image pair was acquired using Google’s similar image search function, for a total of 111 image pairs (222 images). 21 image pairs were used for practice, and the remaining 90 were used during the in-scan encoding and retrieval tasks (see Supplementary Figure 5 for example image pairs). Each image was paired with a short descriptive audio title in a synthesized female voice (https://neospeech.com; voice: Kate) during encoding runs; this title served as a retrieval cue during the in-scan retrieval task. Two videos used for model training (720 by 480 pixels; 10m 25s and 10m 35s in length) were composed of a series of short (~4s) clips drawn from YouTube and Vimeo, containing a wide variety of themes (e.g. still photos of bugs, people performing manual tasks, animated text, etc.). One additional video cut from “Indiana Jones: Raiders of the Lost Ark” (1024 by 435 pixels; 10m 6s in length) was displayed while in the scanner, but the associated data was not used in this experiment.

### Procedure

Before undergoing MRI, participants were trained on a practice version of the task incorporating 21 practice image pairs. Inside the MRI scanner, participants completed three video viewing runs and three encoding-retrieval sets. The order of the runs was as follows: first video viewing run (short clips 1), second video viewing run (short clips 2), third video viewing run (Indiana Jones clip), first encoding-retrieval set, second encoding-retrieval set, third encoding-retrieval set. A high-resolution structural scan was acquired between the second and third encoding-retrieval sets, providing a break.

Video viewing runs were 10m 57s seconds long. For each run, participants were instructed to pay attention while the video (with audio) played within the center of the screen. The order of the videos was the same for all participants.

Encoding-retrieval sets were composed of one encoding run followed by one retrieval run. Each set required the participants to first memorize and then recall 30 images drawn from 30 image pairs. The image pairs within each set were selected randomly, with the constraint that no image pair could be used in more than one set. The image selected from each image pair to be presented during encoding was counterbalanced across subjects. This experimental procedure was designed to limit the concurrent memory load to 30 images.

Encoding runs were 6m 24s long. Each run started with 10s during which instructions were displayed on-screen. Trials began with the appearance of an image in the center of the screen (1.8s), accompanied by a simultaneous descriptive audio cue (e.g. a picture depicting toddlers would be coupled with the spoken word “toddlers”). Images occupied 800 by 600 pixels of a 1024 by 768 pixel screen. Between trials, a cross-hair appeared in the center of the screen (font size = 50) for 1.7s. Participants were instructed to pay attention to each image and to encode as many details as possible so that they could visualize the images as precisely as possible during the imagery task. To assess ongoing engagement with the task, the participants also performed a 1-back task requiring the participants to press “1” if the displayed image was the same as the preceding image, and “2” otherwise. Within each run, stimuli for the 1-back task were randomly sampled with the following constraints: 1) each image was repeated exactly four times in the run (120 trials per run; 360 for the entire scan), 2) there was only one immediate repetition per image, and 3) the other two repetitions were at least 4 items apart in the 1-back sequence.

Retrieval runs were 9m 32s long. Each run started with 10s during which instructions were displayed on-screen. Thirty images were then cued once each (the order was randomized), for a total of 30 trials per run (90 for the entire scan). Trials began with an image title appeared in the center of the screen for 1s (font = Courier New, font size = 30). After 1s, the title was replaced by an empty rectangular box shown in the center of the screen (6s), and whose edges corresponded to the edges of the stimulus images (800 by 600 pixels). Participants were instructed to visualize the image that corresponded to the title as accurately and in as much detail as they could within the confines of the box. Once the box disappeared, participants were prompted to rate the vividness (defined as the relative number of recalled visual details specific to the cued image presented during encoding) of their mental image on a 1-4 scale (3s) using a four-button fiber optic response box (right hand; 1 = right index finger; 4 = right little finger). This was followed by the appearance of a probe image (800 by 600 pixels) in the center of the screen (3s), that was either the same as or similar to the trial’s cued image (i.e. either the image shown during encoding or its pair). While the image remained on the screen, the participants were instructed to respond with “1” if they thought that the image was the one seen during encoding (old), or “2” if the image was new (responses made using the response box). Following the disappearance of the image, participants were prompted to rate their confidence in their old/new response on a 1-4 scale (2s) using the response box. Between each trial, a cross-hair (font size = 50) appeared in the center of the screen for either 1, 2 or 3 seconds.

Randomization sequences were generated such that both images within each image pair (image A and B) were presented equally often during the encoding runs across subjects. During retrieval runs each image appeared equally often as a matching (encode A -> probe A) or mismatching (encode A -> probe B) image across subjects. Due to the need to remove several subjects from the analyses, stimulus versions were approximately balanced over subjects.

### Setup and Data Acquisition

Participants were scanned with a 3.0-T Siemens MAGNETOM Trio MRI scanner using a 32-channel head coil system. A high-resolution gradient-echo multi-slice T1-weighted scan coplanar with the echo-planar imaging scans (EPIs) was first acquired for localization. Functional images were acquired using a multiband EPI sequence sensitive to BOLD contrast (22 × 22 cm field of view with a 110 × 110 matrix size, resulting in an in-plane resolution of 2 × 2 mm for each of 63 2-mm axial slices; repetition time = 1.77 sec; echo time = 30ms; flip angle = 62 degrees). A high-resolution whole-brain magnetization prepared rapid gradient echo (MP-RAGE) 3-D T1 weighted scan (160 slices of 1mm thickness, 19.2 × 25.6 cm field of view) was also acquired for anatomical localization.

The experiment was programmed with the E-Prime 2.0.10.353 software (Psychology Software Tools, Pittsburgh, PA). Visual stimuli were projected onto a screen behind the scanner made visible to the participant through a mirror mounted on the head coil.

### fMRI Preprocessing

Functional images were converted into NIFTI-1 format, motion-corrected and realigned to the average image of the first run with AFNI’s (Cox 1996) *3dvolreg* program. The maximum displacement for each EPI image relative to the reference image was recorded and assessed for head motion. The average EPI image was then co-registered to the high-resolution T1-weighted MP-RAGE structural using the AFNI program align_epi_anat.py (Saad et al, 2009).

The volumetric functional data for each experimental task (video viewing, 1-back encoding task, retrieval task) was then projected to a subject-specific cortical surface generated by Freesurfer 5.3 (Dale, Fischl & Sereno, 1999). The target surface was a spherically normalized mesh with 32000 vertices that was standardized using the resampling procedure implemented in the AFNI program *MapIcosahedron* (Argall, Saad & Beauchamp, 2006). To project volumetric imaging data to the cortical surface we used the AFNI program *3dVol2Surf* with the “average” mapping algorithm, which approximates the value at each surface vertex as the average value among the set of voxels that intersect a line along the surface normal connecting the white matter and pial surfaces.

The three video scans (experimental runs 1-3), because they involved a continuous stimulation paradigm, were directly mapped to the surface without any pre-processing to the cortical surface. The three retrieval scans (runs 5, 7, 9) were first divided into a sequence of experimental trials with each trial beginning (t=−2) two seconds prior to the onset of the retrieval cue (verbal label) and ending 32 seconds later in two second increments. These trials were then concatenated in time to form a series of 90 trial-specific time-series, each of which consisted of 16 samples. The resulting trial-wise data blocks were then projected onto the cortical surface. To facilitate separate analyses of the “imagery” and “old/new judgment” retrieval data, a regression approach was implemented. For each trial, the expected hemodynamic response associated with each task was generated by convolving a series of instantaneous impulses over the task period (10 per second; imagery: 61; old/new: 31) with the SPM canonical hemodynamic response. Estimates of beta coefficients for each trial and task were computed via a separate linear regression per trial (each with 16 samples: one per time point), with vertex activity as the dependent variable, and the expected hemodynamic response values for the “recall” and “old/new judgment” tasks as independent variables. Finally, data from the three encoding scans (runs 4, 6, 8) were first analyzed in volumetric space using a trial-wise regression approach, where the onset of each image stimulus was modelled with a separate regressor formed from a convolution of the instantaneous impulse with the SPM canonical hemodynamic response. Estimates of trial-wise beta coefficients were then computed using the “least squares sum” (Mumford, Turner, Ashby & Poldrack, 2012) regularized regression approach as implemented in the AFNI program *3dLSS*. The 360 (30 unique images per run, 4 repetitions per run, 3 total runs) estimated beta coefficients were then projected onto the cortical surface with *3dVol2Surf*.

### Deep Neural Network Image Features

We used the TensorFlow implementation of the VGG16 deep neural network (DNN) model (Simonyan & Zisserman, 2014; see http://www.cs.toronto.edu/~frossard/post/vgg16 for the implementation used). The VGG16 model consists of a total of thirteen convolutional layers and three fully connected layers. 90 image pairs from the memory task and 3775 video frames (3 frames per second; taken from the two short-clip videos; video 1: 1875 frames; video 2: 1900 frames) were resized to 224 × 224 pixels to compute outputs of the VGG16 model for each image/frame. The outputs from the units in the second convolutional layer (layer 2), the seventh convolutional layer (layer 7), the last convolutional layer (layer 13), and the final fully connected layer (layer 16) were treated as vectors corresponding to low-level visual features, mid-level visual features, high-level visual features and semantic features, respectively. Layer selection was performed manually by inspection of filter activations (see Figure 1c for example activations). To account for the low retinotopic spatial resolution resulting from participants eye movements, the spatial resolution of the convolutional layers (the fully connected layer has no explicit spatial representation) was reduced to 3 by 3 (original resolution for layer 2: 224 by 224; layer 7: 56 by 56; layer 13: 14 by 14). The resultant vector length of low-level visual features, mid-level visual features, high-level visual features and semantic features was 576, 2304, 4608 and 1000, respectively. Convolutional layer activations were log-transformed to improve prediction accuracy (Naselaris et al., 2015).

### Encoding Model

An encoding model was estimated for each of the four feature levels and each individual voxel (Naselaris et al., 2015). Let *v*_*it*_ be the signal from voxel *i* during trial *t*. The encoding model for this voxel is:

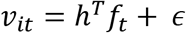

Here *f*_*t*_ is a 100 × 1 vector of 100 image features associated with the current trial/image (only the 100 features with the largest positive correlations with voxel activity were selected to make the computation tractable), *h* is a 100 × 1 vector of model parameters that indicate the voxel’s sensitivity to a particular feature (the superscript *T* indicates transposition) and ϵ is zero-mean Gaussian additive noise.

The model parameters *h* were fit using non-negative lasso regression (R package “nnlasso”; Mandal & Ma, 2016) trained on data drawn from the encoding and movie viewing (excluding the Indiana Jones video) tasks using 3-fold cross validation over the encoding data (all movie data was used in each fold). The non-negative constraint was included to reduce the possibility that a complex linear combination of low-level features may approximate one or more high-level features. The regularization parameter (lambda) was determined by testing 5 log-spaced values from approximately 1/10000 to 1 (using the nnlasso function’s path feature). For each value of the regularization parameter, the model parameters *h* were estimated for each voxel and then prediction accuracy (sum of squared errors; SSE) of the recognition data was measured using 3-fold cross validation. For each voxel, the model parameters *h* that yielded the highest prediction accuracy were retained for image decoding.

### Image Decoding

Encoding models were used to predict neural activity during recall for each unique combination of subject, feature-level, ROI, and retrieval trial (74 bilateral cortical FreeSurfer ROIs). The accuracy of this prediction was assessed as follows: 1) for each combination of subject, feature-level, and ROI the predicted neural activation patterns for the 90 images viewed during the encoding task were generated using a model that was trained on the movie and encoding task data, excluding data from encoding trials wherein the predicted image was viewed using 3-fold cross validation. 2) for each retrieval trial, the predictions were correlated (across vertices within the given ROI) with the observed neural activity during recall resulting in 90 correlation coefficients. 3) the correlation coefficients were ranked in descending order, and the rank of the prediction associated with the recalled image was recorded (1 = highest accuracy, 90 = lowest accuracy). 4) this rank was then subtracted from the mean rank (45.5) so that 0 was chance, and a positive value indicated greater-than-chance accuracy for the given trial (44.5 = highest accuracy, −44.5 = lowest accuracy).

### Seed ROI Selection

Separate ROIs were selected from all Freesurfer ROIs for each of the four feature levels. The procedure for generating weight values for each ROI (Figure 3a) was as follows: 1) get the average classification accuracy across subjects during image perception (data taken from the old/new recognition task during the retrieval blocks) for each feature-level and ROI. 2) z-score classification accuracy across ROIs for each feature-level. 3) set all values less than zero to zero. 4) for each ROI and feature-level, subtract the greatest value associated with the other feature-levels from the target feature’s value. 5) set all values less than zero to zero. 6) normalize the values across ROIs to sum up to one (i.e. divide each value by the sum of all values) for each feature level. 7) set values less than .05 to 0 to retain only those ROIs that have a strong association with the feature-level. 8) normalize the values across ROIs for each feature level.

### Feature-Specific Informational Connectivity

For the FSIC analyses, correlations of image classification accuracy (rank measure) during imagery were performed across ROIs. Separate correlations were performed for all combinations of four seeds (selected from all Freesurfer ROIs as outlined in “Seed ROI Selection”) and all other ROIs not included within the seed. For each seed, classification accuracy was drawn from the seed’s associated feature level. The correlations were calculated with a linear mixed-effects (LME) model on data from all episodic recall trials, wherein classification accuracy for the seed ROI was the dependent variable (DV), classification accuracy for each of the four feature levels within the target ROI were the independent variables (IV), and participant and image were crossed random effects (random-intercept only, due to model complexity limitations). Statistical assessments were performed using bootstrap analyses, calculated with the BootMer function (Bates et al. 2015) using 1000 samples and corrected for multiple comparisons across ROIs using FDR (Benjamini & Hochberg, 1995).

### fMRI Data Simulation

The simulation used the same experimental structure and stimuli (for training and testing the models) as the true experiment. For each simulated subject, 800 artificial voxels were created, with each voxel containing one, randomly selected, feature extracted from the CNN VGG16 as described in the “Deep Neural Network Image Features” section. For each voxel, the feature-specific activation associated with the video frame or image presented at each time-point or trial was used to simulate the voxel’s activity. Voxels were grouped into 8 ROIs with 100 voxels each. There were 2 ROIs per feature-level (one representing the seed ROI, and the other representing the target ROI), such that features assigned to the voxels in each ROI were extracted from the assigned level. The two ROIs assigned the same level contained identical features, i.e. they were duplicates, except for the subsequent application of independent gaussian noise. For the analysis depicted in Supplementary Figure 2b, the seed ROIs contained 25 voxels representing each of the four feature-levels (for 100 voxels total). Memory loss during recall was simulated by randomly setting a fraction of the features to zero. The same features were set to zero across ROIs representing the same feature level for a given trial, simulating cross-ROI information transfer. Trial-by-trial variation in memory accuracy was simulated by varying the fraction of feature loss over trials (randomly selected using a uniform distribution from 40 – 95%). Lastly, independent gaussian noise (mean 0, standard deviation 1) was added to all data, with the SNR varying across simulated subjects (either 15, 25 or 35%, equally distributed), to simulate all unaccounted-for variation in voxel activity, and individual variations thereof.

### Linear Models and Statistics

Statistical assessment of mean neural reactivation (Figure 2 and 3a) was performed using a separate LME model for each ROI, with neural reactivation as the DV and subject and image as crossed random effects. Confidence intervals and p-values were calculated with bootstrap statistical analyses (1000 samples) using the BootMer function (Bates et al. 2015) and corrected for multiple comparisons across ROIs using false discovery rate (FDR; Benjamini & Hochberg, 1995). For the within-subject correlations between feature-specific reactivation, vividness ratings (Figure 5b), and recognition accuracy, LME models were used, with vividness ratings or recognition accuracy (correct vs incorrect) as the dependent variable (DV), the four neural reactivation measures for each combination of ROI (lower-level and higher-level) and feature-level (lower-level and higher-level) as independent variables (IV), and participant and image as crossed random effects (random-intercept only, due to model complexity limitations). Confidence intervals and p-values were calculated with bootstrap statistical analyses (1000 samples) using the BootMer function and corrected for multiple comparisons across coefficients using FDR. For the between-subject correlations between feature-specific reactivation and recognition accuracy (Figure 5c), a single linear model was used, with recognition accuracy (correct vs incorrect) as the dependent variable (DV) and the four neural reactivation measures as independent variables (IV). Confidence intervals and p-values were generated with bootstrap statistical analyses (1000 samples) and corrected for multiple comparisons using FDR across coefficients—including the four coefficients from the within-subject recognition accuracy LME (i.e. eight coefficients in total).

## Author Contributions

Conceptualization, M.B.B., B.R.B.; Methodology, M.B.B., B.R.B., F.A.; Software, M.B.B., B.R.B.; Formal Analysis, M.B.B.; Investigation, F.A.; Data Curation, M.B.B., B.R.B.; Writing – Original Draft, M.B.B.; Writing – Review and Editing, M.B.B., B.R.B.; Visualization, M.B.B.; Supervision, B.R.B.

