## Supplementary Figure 1 for "Feature-Specific Neural Reactivation during Episodic Memory"

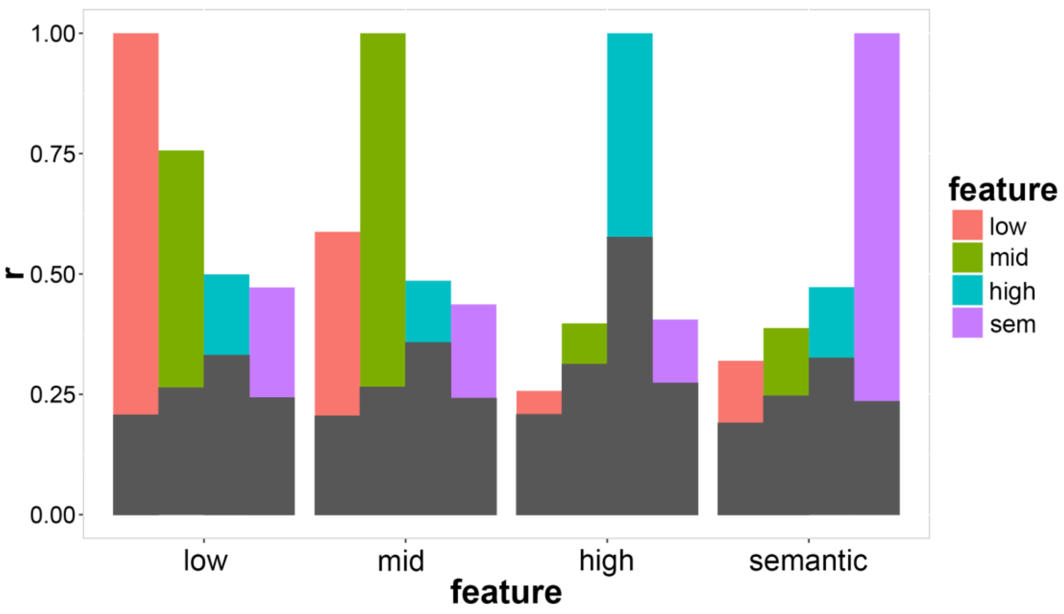


**Supplementary Figure 1. Correlation Between Features from Different Layers of a CNN.** Grey bars indicate 95% upper bounds of the null distribution (1000 sample permutation). All cross-feature-level correlations were found to be significantly greater than chance, with the correlation magnitude inversely proportional to the distance between the layers. To generate the correlation values, activation vectors (with one value for all 90 image pairs, e.g. 180 images/elements in length) for each of *M* node/features in one layer (represented by the x-axis) were correlated with all *N* node/features in another layer (represented by the legend/color), producing an $M \times N$ correlation matrix. For each row of the correlation matrix, the maximum correlation value was extracted, and the resulting *M* correlation values were averaged (mean).
