## Supplementary Figure 2 for "Feature-Specific Neural Reactivation during Episodic Memory"

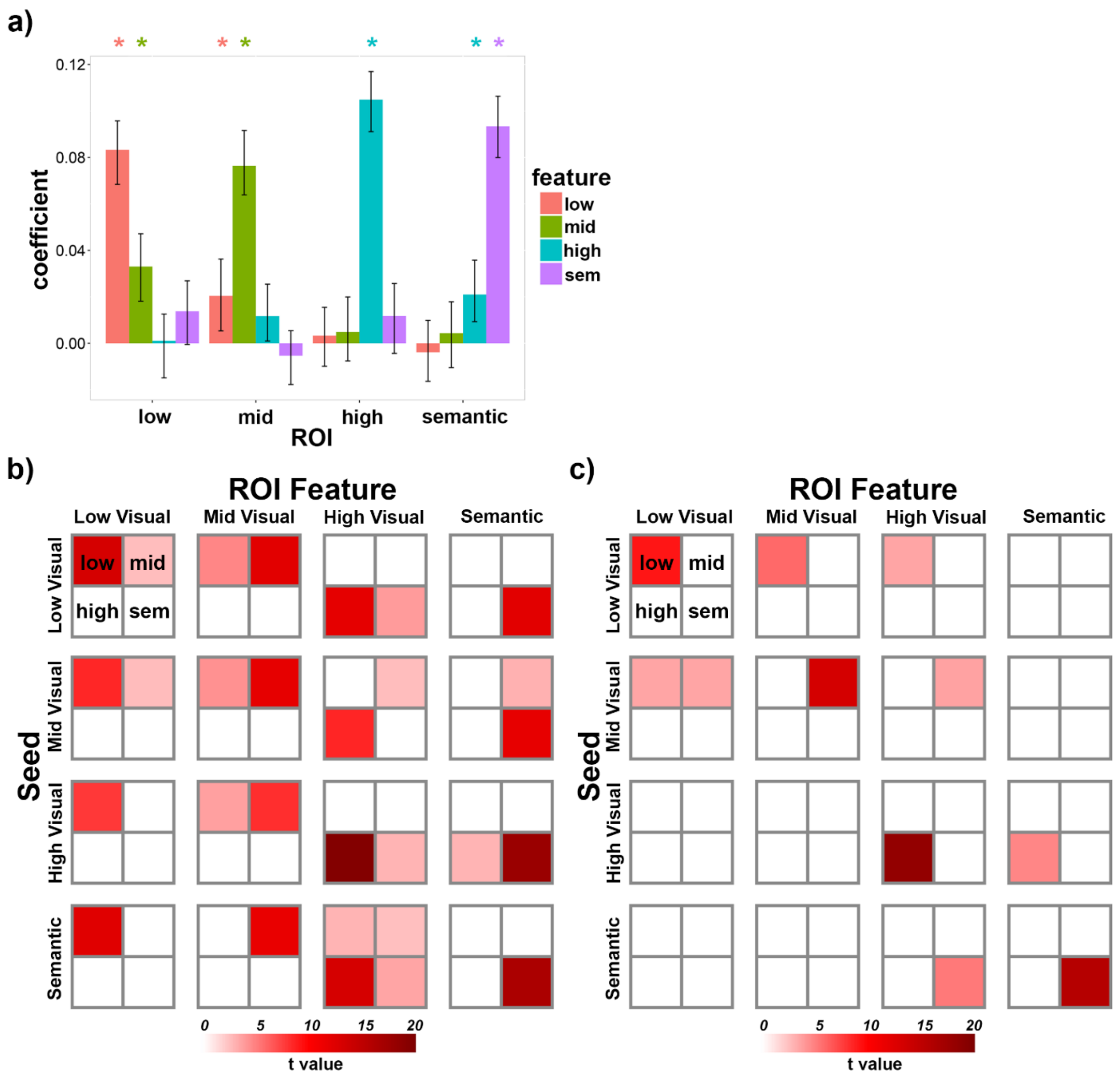


**Supplementary Figure 2. Simulated Results for Feature-Specific Informational Connectivity.** fMRI data was simulated (200 simulated subjects; see Methods section) and then run through the processing pipeline for FSIC (see Methods section) to validate the approach. ROIs only contain features from the indicated feature-level. a) FSIC results for all combinations of ROI and feature-level, assuming identical trial-by-trial memory accuracy across feature-levels. A separate seed was used for each feature-level, with each containing an equal number of voxels (25) per feature-level. Error bars are 90% CIs; * indicates $p < 0.05$, one-tailed, FDR corrected. The similarity of the results to Figure 3b indicates that the feature specificity of FSIC does not depend on the seed disproportionately representing the target feature-level. c) and d) FSIC results for all combinations of seed ROI/feature-level (rows) and target ROI feature-level (columns), assuming c) identical and d) independent trial-by-trial memory accuracy across feature-levels. The squares divided into four sub-squares represent a simulated brain composed of four target ROIs. Each ROI contains features from one feature-level, as indicated in the top-left corner. t-values are thresholded at $p < 0.05$, one-tailed, FDR corrected. Under the cross-feature-level memory dependence assumption, seed selection had little effect on the results, whereas under the independence assumption, significant effects were primarily limited to the diagonal (i.e. when the seed and the target feature correspond).
