## Supplementary Figure 3 for "Feature-Specific Neural Reactivation during Episodic Memory"

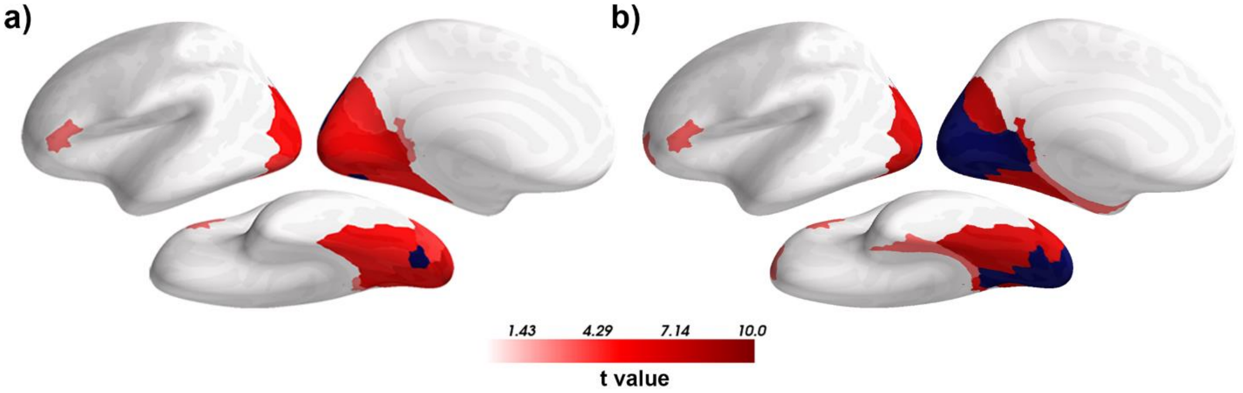


**Supplementary Figure 3. Effect of Seed Size on Mid-Level Feature-Specific Informational Connectivity.** a) FSIC using the mid-level seed, as depicted in Figure 4. b) FSIC using a combination of the low- and mid-level seeds, as depicted in Figure 5a: "low + mid". t-values are thresholded at $p < 0.05$, one-tailed, FDR corrected. Results are nearly identical, indicating that the difference in the extent of low- and mid-level neural reactivation was not due to the relatively small size of the mid-level ROI.
