## Supplementary Figure 4 for "Feature-Specific Neural Reactivation during Episodic Memory"

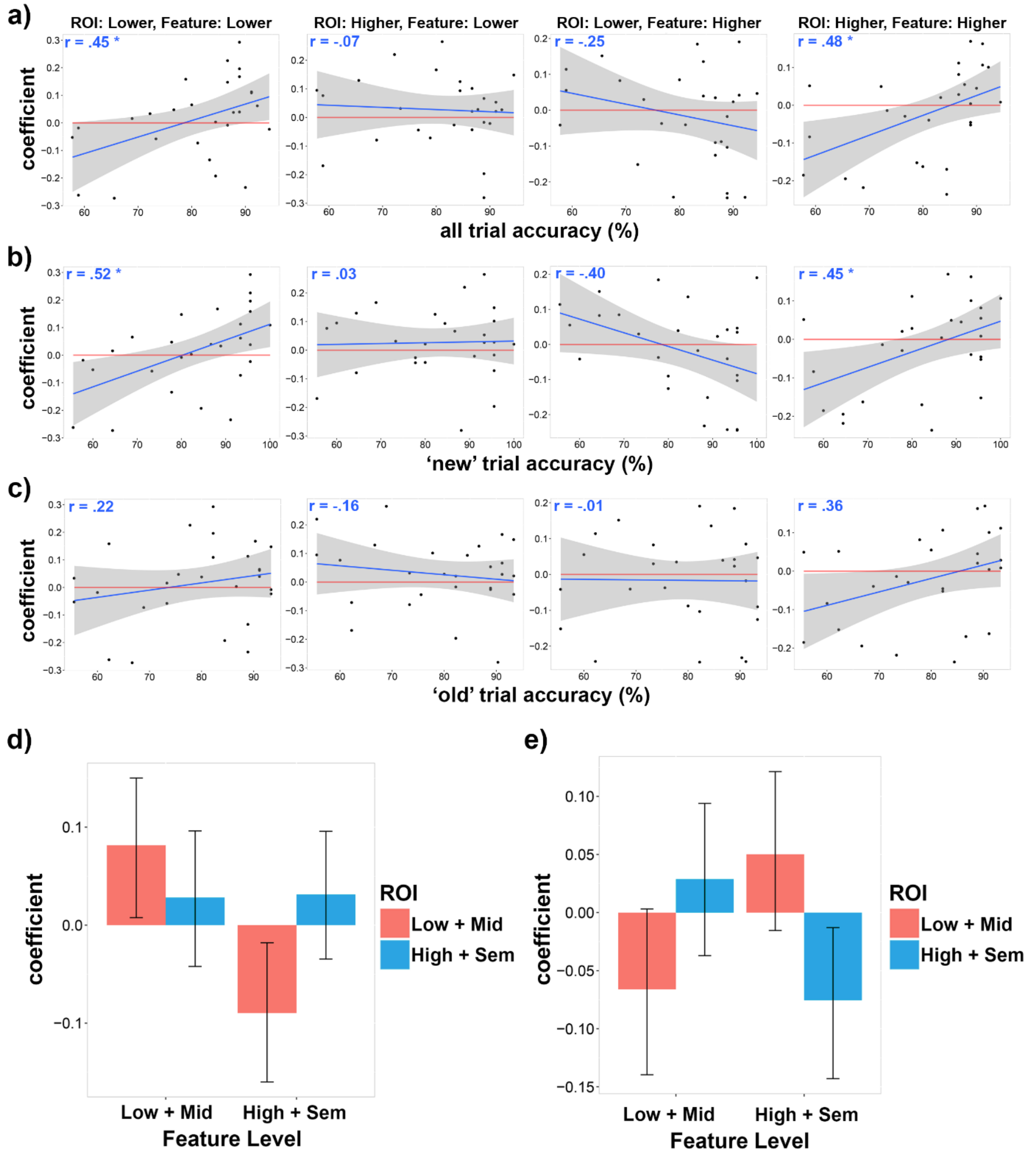


**Supplementary Figure 4. Individual Differences in the Correlation Between Recognition Accuracy and Neural Reactivation during Recall.** a-c) The between-subject correlation between average old/new task accuracy (percentage correct for a) all trials, b) 'new'/lure trials, and c) 'old' trials) and the within-subject partial correlation coefficient between old/new accuracy and neural reinstatement for all four combinations of ROI and feature-level. Points represent subjects; grey region indicates 95% CI; * indicates p < .05, two-tailed, FDR corrected.

d-e) Within-subject partial correlations between neural reactivation and old/new task accuracy for all combinations of feature-level and ROI, with the participants divided into two groups: d) the thirteen subjects with the highest average ‘new’/lure trial accuracy, and c) the thirteen subjects with the lowest average ‘new’/lure trial accuracy. Error bars are 95% CI; * indicates $p < .05$, two-tailed, FDR corrected. Consistent with the between-subject results, reactivation encoding lower-level features positively correlated with accuracy (before considering the other coefficients), but only for the high-lure-accuracy group, i.e. subjects who were less likely to label a similar new image as previously seen. When considering all coefficients, no coefficient was significantly greater than zero after correcting for multiple comparisons (FDR).
