## Supplementary Figure 5 for "Feature-Specific Neural Reactivation during Episodic Memory"

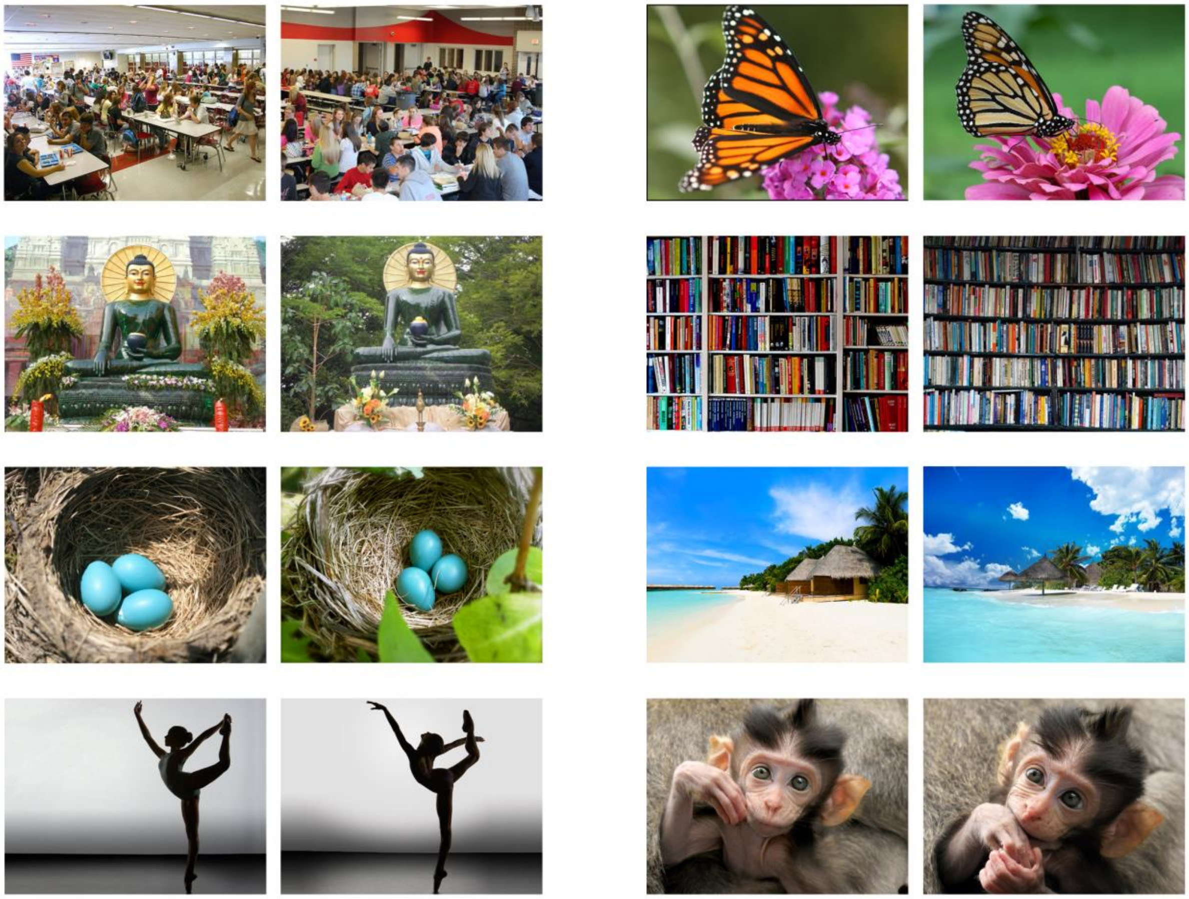

**Supplementary Figure 5. Example of Image Pairs.** Eight randomly selected image pairs out of the ninety total.
